# TDCS MODULATES WORKING MEMORY MAINTENANCE PROCESSES IN HEALTHY INDIVIDUALS

**DOI:** 10.1101/2022.11.23.517774

**Authors:** Stevan Nikolin, Donel Martin, Colleen K. Loo, Tjeerd W. Boonstra

## Abstract

**Background:** The effects of tDCS at the prefrontal cortex are often investigated using cognitive paradigms, particularly working memory tasks. However, the neural basis for the neuromodulatory cognitive effects of tDCS, including which subprocesses are affected by stimulation, is not completely understood.

**Aims:** We investigated the effects of tDCS on working memory task-related spectral activity during and after tDCS to gain better insights into the neurophysiological changes associated with stimulation.

**Methods:** We reanalysed data from 100 healthy participants grouped by allocation to receive either Sham (0 mA, 0.016 mA, and 0.034 mA) or Active (1 mA or 2 mA) stimulation during a 3-back task. Electroencephalography (EEG) data was used to analyse event-related spectral power in frequency bands associated with working memory performance.

**Results:** Frontal theta event-related synchronisation (ERS) was significantly reduced post-tDCS in the active group. Participants receiving active tDCS had slower response times following tDCS compared to Sham, suggesting interference with practice effects associated with task repetition. Theta ERS was not significantly correlated with response times or accuracy.

**Conclusions:** tDCS reduced frontal theta ERS post-stimulation, suggesting a selective disruption to working memory cognitive control and maintenance processes. These findings suggest that tDCS selectively affects specific subprocesses during working memory, which may explain heterogenous behavioural effects.

## Introduction

Transcranial direct current stimulation (tDCS), a form of non-invasive brain stimulation capable of modulating cortical activity (Jog et al., 2016; Wiesman et al., 2018), is increasingly used as a technique to investigate cognitive functioning across multiple domains (Berryhill & Martin, 2018; Berryhill, Peterson, Jones, & Stephens, 2014). Of these, one of the most researched is working memory. Working memory has been described as a ‘cognitive primitive’ due to its central role as a core executive function (Diamond, 2013; Pennington, 1994). It is a complex short-term memory storage system distinguished by its capacity to maintain and manipulate a limited amount of temporally ordered information (Aben, Stapert, & Blokland, 2012; A. Baddeley, 2003; A. D. Baddeley & Hitch, 1974). Meta-analyses have demonstrated a significant effect of tDCS on working memory. Specifically, healthy participants were found to improve on measures of response time, whereas accuracy was increased in neuropsychiatric populations (Brunoni & Vanderhasselt, 2014; Dedoncker, Brunoni, Baeken, & Vanderhasselt, 2016; Hill, Fitzgerald, & Hoy, 2016; Lee, Lee, & Kang, 2021), although some of these findings have been questioned (Medina & Cason, 2017).

Working memory requires the coordinated activity of a network of brain regions (D’Esposito, 2007), including the dorsolateral prefrontal cortex (DLPFC), a key node within the frontoparietal cognitive control network (Owen, McMillan, Laird, & Bullmore, 2005). The cortical dynamics supporting working memory may be captured by measuring task-induced neural oscillations using electroencephalography (EEG) (Deiber et al., 2007); for a review see Pavlov and Kotchoubey (2020). Briefly, maintenance processes are reflected in frontal theta (4-8 Hz) activity (Brzezicka et al., 2019; Fernández, Pinal, Díaz, & Zurrón, 2021) and incorporate the DLPFC, occipitotemporal, and other brain regions (Gazzaley, Rissman, & D’esposito, 2004). In particular, event-related synchronisation (ERS) in the theta band is associated with cognitive control efforts during working memory processing (Cavanagh & Frank, 2014; Duprez, Gulbinaite, & Cohen, 2020), and positively correlates with cognitive load (Jensen & Tesche, 2002; Pesonen, Hämäläinen, & Krause, 2007). Alpha frequency (8-12Hz) event-related desynchronization (ERD), signifying a reduction in spectral power, has been associated with attentional processes (Fodor, Marosi, Tombor, & Csukly, 2020; Hanslmayr, Gross, Klimesch, & Shapiro, 2011), including visual processing (Wianda & Ross, 2019) and top-down executive control thought to protect working memory maintenance from external distractors (Bonnefond & Jensen, 2012). Beta (13-30 Hz) activity has been linked to active maintenance and item retention for further task requirements (Y. Chen & Huang, 2016; Deiber et al., 2007; Onton, Delorme, & Makeig, 2005). Lastly, gamma ERS has been associated with working memory capacity (C.-M. A. Chen et al., 2014), and increases in working memory effort (Basar-Eroglu et al., 2007). It represents stimulus encoding and propagation of working memory items from sensory regions to those associated with higher-order cortical activity (Novikov & Gutkin, 2018). Investigating task-induced spectral changes may hence provide insights into the mechanism of action by which tDCS modulates working memory processes.

In comparison with the behavioural effects of tDCS on working memory, relatively few studies have explored tDCS-induced changes using EEG measures of cortical dynamics. Gamma ERS has been shown to increase during 2- and 3-back working memory tasks following prefrontal tDCS (Boudewyn, Roberts, Mizrak, Ranganath, & Carter, 2019; Hoy, Bailey, Arnold, & Fitzgerald, 2015; Ikeda, Takahashi, Hiraishi, Saito, & Kikuchi, 2019), but others have not observed these effects (Hill, Rogasch, Fitzgerald, & Hoy, 2018; Murphy et al., 2020). Theta ERS acquired during a 2-back task has been shown to increase following stimulation as compared to sham (Hoy et al., 2013), and to cathodal tDCS (Zaehle, Sandmann, Thorne, Jäncke, & Herrmann, 2011). These studies also report conflicting findings for the alpha frequency band; while Zaehle et al. (2011) found an attenuation of ERD (i.e., a relative increase of spectral power) following anodal prefrontal tDCS relative to cathodal stimulation, Hoy et al. (2013) reported greater ERD as compared to sham. Collectively, these findings imply beneficial effects of tDCS to maintenance operations and encoding of stimuli to working memory, as reflected by greater theta and gamma ERS, respectively. However, due to the mixed nature of findings in the literature, the impact of tDCS on cognitive processes associated with working memory requires further elucidation.

Therefore, we investigated the neuromodulatory effects of tDCS on ERS/ERD during a working memory task paradigm in a large sample of healthy participants (n=100). We aimed to determine which frequency-specific cognitive subprocesses are modulated by tDCS. Working memory performance outcomes and EEG-based time-frequency measures were assessed in the period immediately following stimulation. Furthermore, we sought to investigate changes to cortical dynamics during tDCS as the concurrent effects of tDCS have been minimally investigated using EEG due to the inherent difficulties posed by stimulation artefacts.

## Methods

### Participants

Detailed methods and primary behavioural results are reported in Nikolin, Martin, Loo, and Boonstra (2018). The current study involves a reanalysis of EEG and behavioural data. Briefly, a total of 100 participants (age: 22.9 ± 4.3 years: males: 47; females: 53) were recruited. Participant exclusion criteria included significant psychological or neurological illness, excessive alcohol or illicit substance abuse, smoking, and ambidextrous or left-handed volunteers assessed using the Edinburgh handedness test (Oldfield, 1971). Participants were evenly allocated to one of five conditions in a parallel group, single blind design. To leverage the large dataset available to investigate working memory effects of tDCS, participants receiving Sham1, Sham2, and Off stimulation were collectively grouped into a Sham category (n = 60), and participants receiving either 1 mA or 2 mA stimulation were likewise categorised as receiving Active stimulation (n = 40). An *a priori* power analysis was not possible given conflicting findings in the literature for tDCS effects on ERS/ERD activity associated with working memory. However, our sample size is sufficient to detect a medium-sized effect (Cohen’s *h* = 0.57) with two-tailed α = 0.05 and 1 - β = 0.80 and is to the best of our knowledge the largest such investigation to date. The study was approved by the University of New South Wales (UNSW) Human Research Ethics Committee (HC13278).

### Procedure

The study consisted of two experiments (Fig. 1), which only differed methodologically in the timing of the baseline 5-min resting-state EEG (results not reported here). Participants were allocated to stimulation conditions in each experiment using identical stratified randomisation methods according to baseline working memory performance (see Nikolin et al. (2018) for stratification thresholds). Of relevance to the current study, participants performed the visual 3-back working memory task, adapted from Mull and Seyal (2001) before, during, and post-tDCS. The 3-back task was administered via Inquisit software (Version 4, Millisecond Software). Participants were presented a series of letters (A - J) each briefly flashing on the screen for 30 ms with an interval of 2000 ms between letter stimuli and were required to press the spacebar on a standard keyboard when the letter presented on the screen matched a letter observed three trials previously. The task was presented for approximately 7 minutes, comprising 40 target stimuli and 180 distractors. Participants were given the opportunity to practise the 3-back task prior to starting the experiment to ensure task instructions were understood. Task performance was assessed using response time (RT) for correct responses and d-prime, a measure of discriminative sensitivity (Haatveit et al., 2010). D-prime was calculated using a z-score transformation of the difference between the percentage of correct responses and incorrect responses (i.e., false alarms).

**Figure 1.**
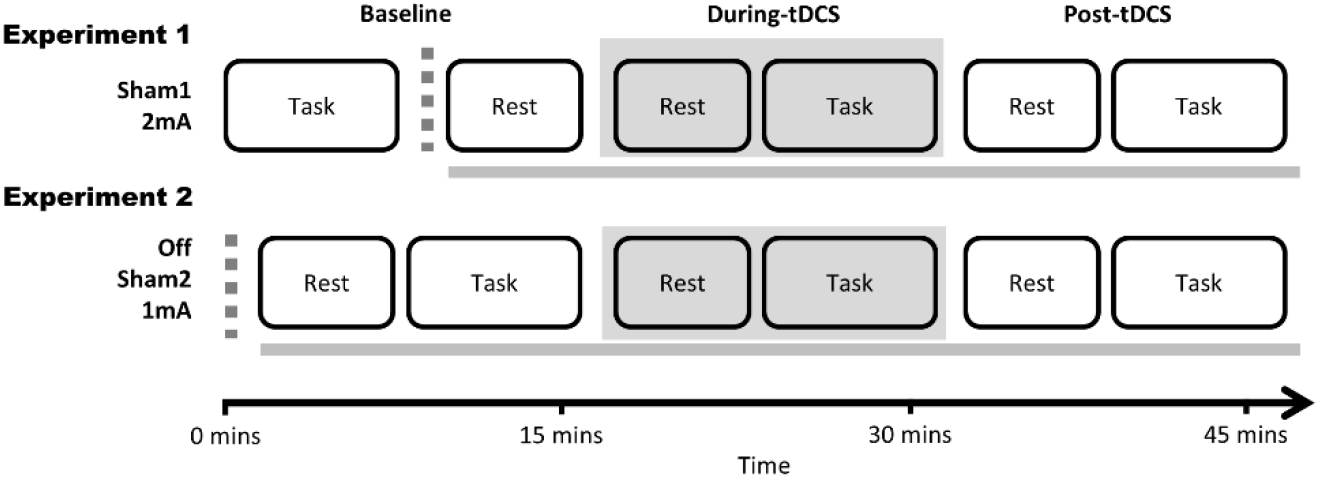
Study protocol. Recruitment occurred in two experiments; in Experiment 1 participants were allocated to receive either Sham1 (0.016mA) or 2mA tDCS; in Experiment 2 participants were allocated to receive either Sham2 (0.034mA), 1mA tDCS, or no stimulation at all in an Off condition (0 mA) in which the electrode leads were left unplugged for the entirety of the experiment. The dotted grey line represents when EEG setup occurred and took approximately 15-20 minutes. The solid grey line shows when EEG was collected.

### Transcranial direct current stimulation

Stimulation was given for 15 min using 4 cm × 4 cm galvanised rubber electrodes (16 cm^2^), with saline soaked sponges placed between the electrodes and the skin to improve conductivity and minimise the risk of skin lesions (Loo et al., 2011). The anode was placed on the left DLPFC (F3 according to the International 10–20 EEG system), a region associated with verbal working memory (Barbey, Koenigs, & Grafman, 2013; Cabeza, Dolcos, Graham, & Nyberg, 2002; Kikyo, Ohki, & Sekihara, 2001; Nyberg et al., 2003) and a common target for non-invasive brain stimulation protocols seeking to modulate working memory performance (Keeser et al., 2011; Teo, Hoy, Daskalakis, & Fitzgerald, 2011; Zaehle et al., 2011). Cathodal inhibitory effects are inconsistently observed in studies of cognition (Jacobson, Koslowsky, & Lavidor, 2012), as such the cathode reference electrode was placed contralaterally on the right DLPFC (F4) in accordance with previous tDCS studies. The two active tDCS conditions were delivered at 1 mA and 2 mA using an Eldith DC-stimulator (NeuroConn GmbH, Germany). Sham tDCS conditions were delivered using either an Eldith DC-stimulator (Sham1), or a tDCS-CT stimulator (Sham2; Soterix Medical Inc., New York). Sham protocols involved an initial ramp up and down of current to elicit paraesthetic sensations and preserve participant blinding. The machine default operation during the off-stimulation mode produced a constant background current of 0.016 mA during Sham1, and 0.034 mA during Sham2. A third, ‘Off’, sham condition was included in which the machine was switched on but the electrode leads were left unplugged (i.e. 0 mA).

### Electroencephalography (EEG) data acquisition

EEG data was acquired using a TMSi Refa amplifier (TMS International, Oldenzaal, Netherlands). The amplifier recorded 24-bit resolution data with no in-built filters, except anti-aliasing. Impedance was kept below 50 kΩ, i.e., <0.5% of the input impedance of the EEG amplifier (100MΩ). A 33-channel head cap with water-based electrodes was used to record 31 EEG channels (see Nikolin et al. (2018)). Sites F3 and F4 were reserved for tDCS-electrode channels, which were secured in place using the EEG cap.

EEG data processing was conducted using custom-developed Matlab scripts (v.2020a; MathWorks) in addition to the Fieldtrip toolbox (Oostenveld, Fries, Maris, & Schoffelen, 2011). EEG data was sampled at 1024Hz and filtered using a second-order bandpass filter (0.5 – 70 Hz) to remove low and high frequency noise generated from head movements and muscle activity, and a Butterworth IIR digital notch filter at 50Hz to remove line noise. The EEG data were epoched into 2 s intervals, beginning 0.5 s before stimulus onset and continuing for 1.5 s after each stimulus presentation. Data were inspected using a semi-automated algorithm to remove epochs containing artefacts. Trials in which any individual channel exceeded amplitudes greater than an absolute z-score of 12 relative to other channels were automatically rejected. Remaining trials were rejected following visual inspection. Independent components analysis (ICA) was then used to remove eye blink and muscle artefacts (Delorme, Sejnowski, & Makeig, 2007; Hyvärinen, Hoyer, & Inki, 2001; Makeig, Bell, Jung, & Sejnowski, 1996) using the default *runica* function implementation in Fieldtrip. Finally, EEG data was re-referenced to the common average reference, which reduces the distortion of measured EEG potentials as compared to a linked mastoids reference (Hu, Lai, Valdes-Sosa, Bringas-Vega, & Yao, 2018).

### Time-frequency analysis

The *n*-back working memory task requires continuous and ongoing maintenance and updating cognitive subprocesses. These underlying working memory processes are therefore activated even during non-target and incorrect trials. For this reason, time-frequency power was calculated in single-trial data and then averaged across all trials, i.e., target letters and distractors combined, using a Hanning taper with a fixed 500 ms time window. Spectral power was calculated for each electrode in a range from 1 – 70Hz and represented as event-related synchronisation (ERS) or desynchronisation (ERD). This shows the relative change in power compared to a reference interval at baseline (−500 to 0 ms before stimulus onset).

### Post-tDCS

Best practice suggests the use of a collapsed localiser blind to group condition to select time intervals for EEG time-frequency and event-related analyses (Cohen, 2014; Luck & Gaspelin, 2017). We therefore computed grand-average time-frequency ERS/ERDs for all participants combined at the post-tDCS timepoint. Time windows containing the largest power changes for frequency bands of interest were selected by visual inspection, allowing the time interval of interest to be defined without bias. Using this methodology, we specified values for frontal theta ERS (4 – 6 Hz; 100 to 1000 ms; EEG channels AFz and Fz – see Fig. 2A), alpha ERD (8 – 13Hz; 230 – 580 ms; EEG channels O1, O2, P7, and P8 – see Fig. 2B), early beta ERD (18 – 24Hz; 130 to 430 ms; EEG channels P7 and P8 – see Fig. 2C), late beta ERS (14 – 20Hz; 600 – 1000 ms; EEG channels O1, Oz, O2, and POz – see Fig. 2D); and gamma ERS (30 – 60Hz; 500 – 1000 ms; EEG channels POz and Pz – see Fig. 2E).

**Figure 2.**
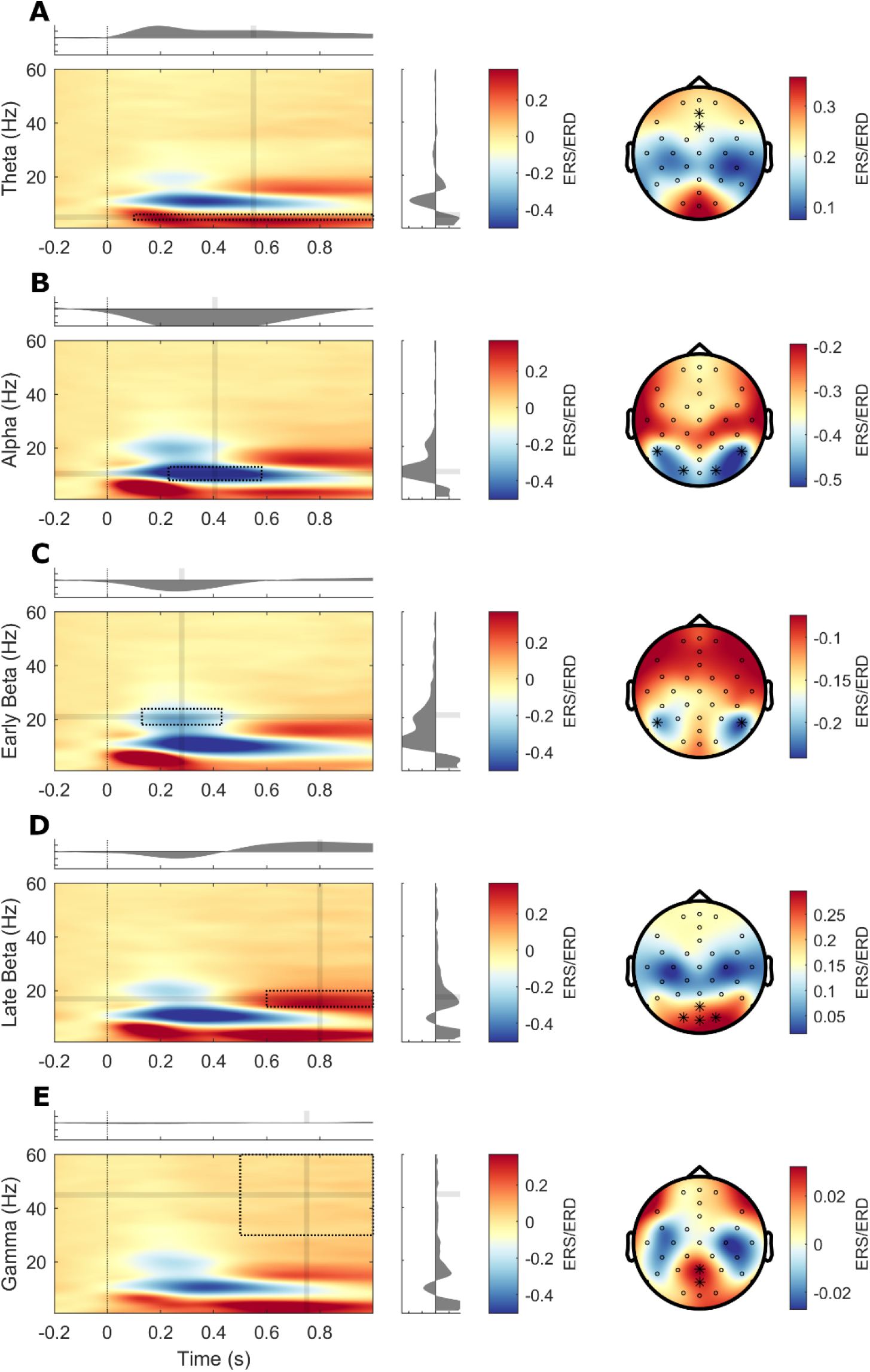
Time-frequency outcomes during *n*-back task. Task-related time-frequency power was generated by averaging brain activity for target and distractor stimuli. Time-frequency plots show event-related synchronisation (ERS) and desynchronization (ERD) at channels highlighted in black for the grand-average of all participants immediately following active tDCS. The black box indicates the time and frequency region of interest. **A)** Frontal theta ERS; **B)** Alpha ERD; **C)** Early beta ERD; **D)** Late beta ERS; **E)** Gamma ERS.

### During-tDCS

The above measures were also analysed during tDCS except for frontal theta, which showed strong signs of contamination from the stimulation artefact. Previous studies have suggested that a high-pass filter is sufficient to generate clean concurrent EEG data during stimulation (Mancini et al., 2015; Marghi et al., 2015), and the online effects of tDCS on EEG activity can be successfully acquired (Giovannella et al., 2018; Schestatsky, Morales-Quezada, & Fregni, 2013; Soekadar, Witkowski, Garcia Cossio, Birbaumer, & Cohen, 2014). Therefore, EEG analyses during tDCS were restricted to alpha frequency bands and higher, effectively adopting a high-pass filter of 8Hz.

### Statistical Analysis

Analyses were performed using SPSS software (IBM SPSS Statistics 26 for Windows, SPSS Inc.). Outliers were identified as those with values greater than three standard deviations from the grand average of all participants and were excluded from the measure/s for which they were an outlier.

### Time-frequency analyses

Due to minor timeline differences in EEG data collection between the two experiment groups, baseline time-frequency measures before tDCS were not available for all participants (i.e., experiment 1). As such, we analysed time-frequency outcomes using separate multivariate analysis of variances (MANOVA) for during- and post-tDCS outcomes, using the independent variable of Condition (Sham: Off, Sham1, and Sham2; and Active: 1 mA and 2 mA). This was done to restrict the number of analyses and so control the Type I error rate, using a significance threshold of *p* < 0.05. A significant MANOVA model (p < 0.05) was followed up with an ANOVA of the significant frequency finding to determine which stimulus conditions (if any) were associated with this difference. Effect sizes were calculated using bootstrapped Cohen’s *d* comparing Active to Sham. Additionally, we performed confirmatory simple univariate linear regressions with time-frequency measures as the dependent variables and the ongoing current intensity participants received over the duration of the stimulation period included as a continuous independent variable (Off: 0 mA; Sham1; 0.016 mA; Sham2: 0.034 mA; and Active conditions of 1 mA and 2 mA).

Follow-up exploratory non-parametric cluster-based permutation tests were performed to identify any significant differences between Sham and Active conditions beyond the *a priori* time interval and channels operationalised for ERS/ERD analyses. This method controls for multiple comparisons while comparing differences across a large spatiotemporal parameter space (Maris & Oostenveld, 2007). Permutation testing was performed across all EEG channels within the time interval 0 – 1000 ms and frequency range of 0.5 – 70 Hz following presentation of *n*-back task stimuli. Participants’ data were permuted 3000 times, and the resulting distributions were compared using independent samples *t*-tests. A value of α < 0.05 was adopted as the two-tailed significance threshold. Due to the limited number of EEG recording channels (i.e., 31) statistically significant clusters were required to comprise only one neighbouring channel.

### Working memory performance

As complete behavioural data was available for all participants before, during, and post-tDCS, we analysed response times for correct responses and d-prime using repeated measures analyses of variance (RMANOVA). Factors included Condition (two levels consisting of Sham: Off, Sham1, and Sham2; and Active: 1 mA and 2 mA), Time (during-tDCS and post-tDCS), and the Condition × Time interaction. Baseline performance was added as a covariate to account for individual differences in working memory. Bootstrapped Cohen’s *d* was used to calculate effect sizes comparing Active to Sham groups during and post-tDCS.

### Correlations

Pearson correlations were used to test the relationship between event-related time-frequency power spectra and working memory performance measures (response times and d-prime) obtained during- and post-tDCS. Correlations were conducted using combined data from all participants included in Active and Sham groups. We controlled for the false discovery rate by applying the Benjamini-Hochberg adjustment for 18 outcomes, setting Q = 0.05 (Benjamini & Hochberg, 1995).

## Results

One participant in the Sham group, allocated to the Off condition, discontinued the experiment during the post-tDCS period. Thus, the sample size in the Sham group was reduced to 59. Table 1 displays baseline demographic information and working memory performance for active and sham groups (independent samples t-tests found no significant between group differences, i.e., all *p*s > 0.05). Side effects were reported in detail in a prior publication (Nikolin et al., 2018). Briefly, all conditions were well tolerated with only minor side effects (e.g., paraesthesia and transient headache) and no serious adverse events.

**Table 1.** Participant characteristics at baseline. Values represent the mean (standard deviation). Response times indicate mean latency for correctly detected target stimuli.

|  | Sham | Active |
| --- | --- | --- |
| Sample (n) | 59 | 40 |
| **Demographic details** |  |  |
| Age | 22.2 (3.1) | 23.9 (5.5) |
| Education (years) | 15.2 (2.2) | 16.0 (2.4) |
| Gender (F/M) | 28/31 | 24/16 |
| **Working Memory** |  |  |
| D-prime | 2.5 (0.7) | 2.6 (0.7) |
| Response time (ms) | 700 (163) | 718 (169) |

### Time-frequency analyses

The number of components rejected during ICA was similar between Sham (5.8; SD = 1.6) and Active participants (5.3; SD = 1.2). Likewise, a similar number of trials were rejected in both groups during-tDCS (Sham: 4.9, SD = 4.5; and Active: 5.4, SD = 4.0) and post-tDCS (Sham: 4.4, SD = 4.7; and Active: 5.8, SD = 5.2).

### During-tDCS

Qualitatively, there were a larger number of outliers identified during-tDCS compared to post-tDCS, with most outliers identified from the Active rather than Sham group. This suggests that stimulation artefacts may be present in the data despite cleaning efforts. Two outliers were excluded for alpha ERD, one for early beta ERD, four for late beta ERS, and lastly nine from gamma ERS analyses. One outlier was identified in the Sham group for gamma ERS.

The MANOVA for time-frequency outcomes extracted during-tDCS was not significant (F = 1.0, *p* = 0.431). Simple univariate linear regression analyses for all time-frequency measures were similarly not significant (all *p*s > 0.05). Cluster-based permutation tests comparing post-tDCS time-frequency ERS/ERD between participants in Active and Sham groups likewise did not reveal significant clusters.

### Post-tDCS

Two outliers were identified and excluded from analysis for late beta ERS and one from gamma ERS in the Active group. For the Sham group, one outlier each was excluded from theta and gamma ERS analyses.

The MANOVA for post-tDCS time-frequency outcomes was significant (F = 2.3, *p* = 0.049). This finding was driven by tDCS effects on frontal theta ERS, in which participants receiving Active tDCS had reduced ERS compared to Sham (F = 4.5; *p* = 0.037; Cohen’s d = -0.42; see Fig 4 and Table 2). Simple univariate linear regression analyses confirmed MANOVA findings, indicating a significant negative relationship between current intensity and frontal theta ERS at the post-tDCS time point (β = -0.06; r^2^ = 0.05; p = 0.029; see appendix Fig. A1). Linear regressions for other time-frequency measures were not significant (all *p*s > 0.05). Cluster-based permutation tests comparing post-tDCS time-frequency ERS/ERD between participants in Active and Sham groups did not identify any significant clusters.

**Table 2.**
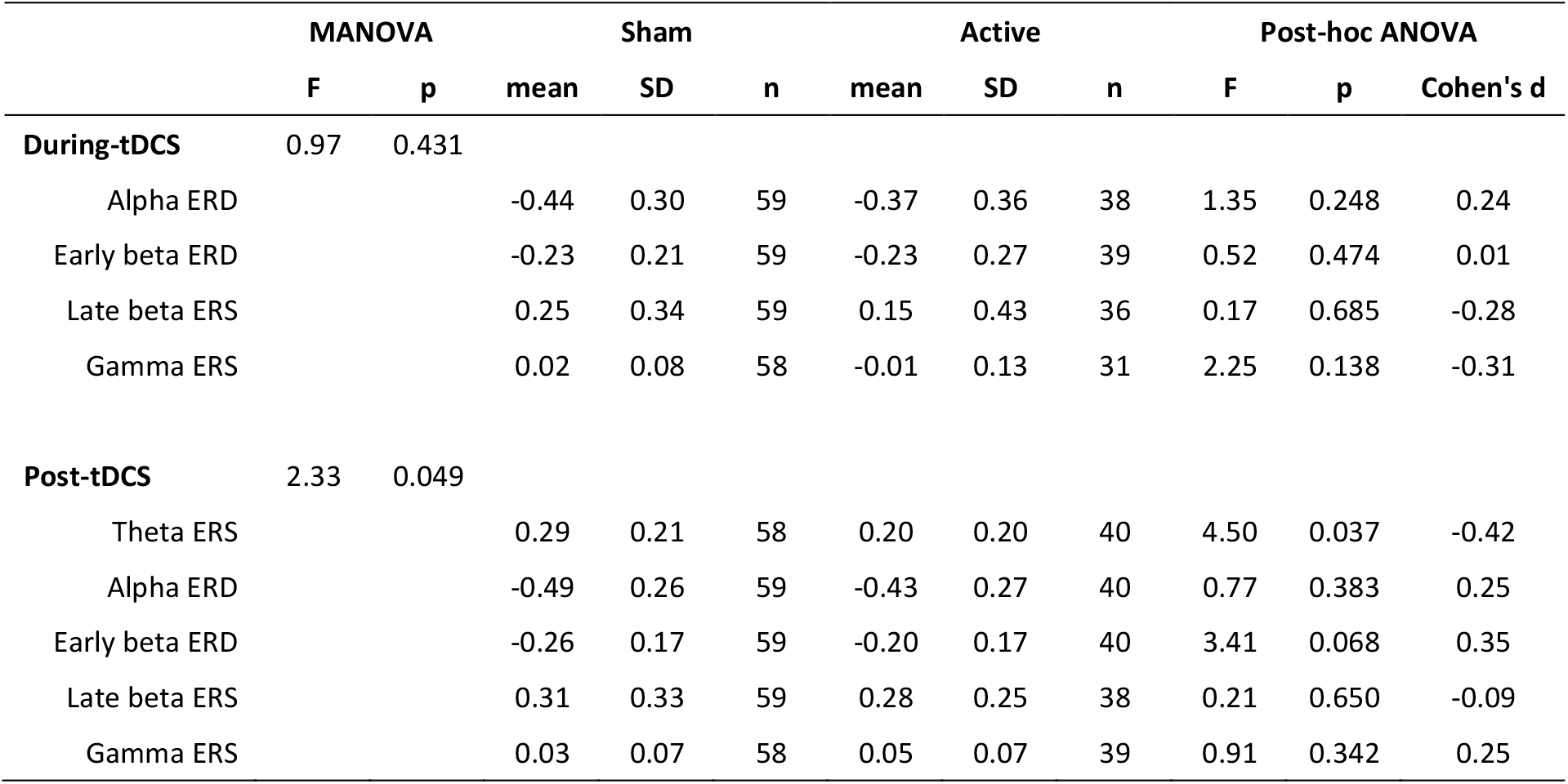
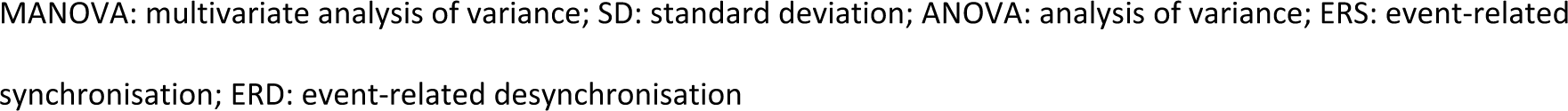
Time-frequency outcomes.

|  | MANOVA |  | Sham |  |  | Active |  |  | Post-hoc ANOVA |  |  |
| --- | --- | --- | --- | --- | --- | --- | --- | --- | --- | --- | --- |
|  | F | p | mean | SD | n | mean | SD | n | F | p | Cohen's d |
| **During-tDCS** | 0.97 | 0.431 |  |  |  |  |  |  |  |  |  |
| Alpha ERD |  |  | -0.44 | 0.30 | 59 | -0.37 | 0.36 | 38 | 1.35 | 0.248 | 0.24 |
| Early beta ERD |  |  | -0.23 | 0.21 | 59 | -0.23 | 0.27 | 39 | 0.52 | 0.474 | 0.01 |
| Late beta ERS |  |  | 0.25 | 0.34 | 59 | 0.15 | 0.43 | 36 | 0.17 | 0.685 | -0.28 |
| Gamma ERS |  |  | 0.02 | 0.08 | 58 | -0.01 | 0.13 | 31 | 2.25 | 0.138 | -0.31 |
| **Post-tDCS** | 2.33 | 0.049 |  |  |  |  |  |  |  |  |  |
| Theta ERS |  |  | 0.29 | 0.21 | 58 | 0.20 | 0.20 | 40 | 4.50 | 0.037 | -0.42 |
| Alpha ERD |  |  | -0.49 | 0.26 | 59 | -0.43 | 0.27 | 40 | 0.77 | 0.383 | 0.25 |
| Early beta ERD |  |  | -0.26 | 0.17 | 59 | -0.20 | 0.17 | 40 | 3.41 | 0.068 | 0.35 |
| Late beta ERS |  |  | 0.31 | 0.33 | 59 | 0.28 | 0.25 | 38 | 0.21 | 0.650 | -0.09 |
| Gamma ERS |  |  | 0.03 | 0.07 | 58 | 0.05 | 0.07 | 39 | 0.91 | 0.342 | 0.25 |

MANOVA: multivariate analysis of variance; SD: standard deviation; ANOVA: analysis of variance; ERS: event-related synchronisation; ERD: event-related desynchronisation

### Working memory performance

Response times on the 3-back task showed a significant main effect of Time (*F* = 7.9, *p* < 0.01, η^2^ = 0.08), indicating shorter latencies post-tDCS relative to during-tDCS. The main effect of Condition was not significant (*F* = 0.4, *p* = 0.53, η^2^ < 0.01; Fig 5), although there was a significant Time × Condition interaction (*F* = 5.8, *p* = 0.02, η^2^ = 0.06). Post-hoc comparisons of the change in response times from during- to post-tDCS revealed a significant difference between Active and Sham (*p* = 0.03, Cohen’s *d* = 0.63) suggesting that response latencies in the Sham group improved at a greater rate in between these time periods as compared to the Active group.

D-prime analysis revealed no significant main effects of Time (*F* = 1.4, *p* = 0.23, η^2^ = 0.02), or Condition (*F* = 1.9, *p* = 0.17, η^2^ = 0.02), and no Time × Condition interaction (*F* = 1.1, *p* = 0.29, η^2^ = 0.01; Fig. 3). Effect sizes comparing Active to Sham participants were small-to-moderate during-tDCS (Cohen’s *d* = -0.38) and post-tDCS (Cohen’s *d* = -0.17).

**Figure 3.**
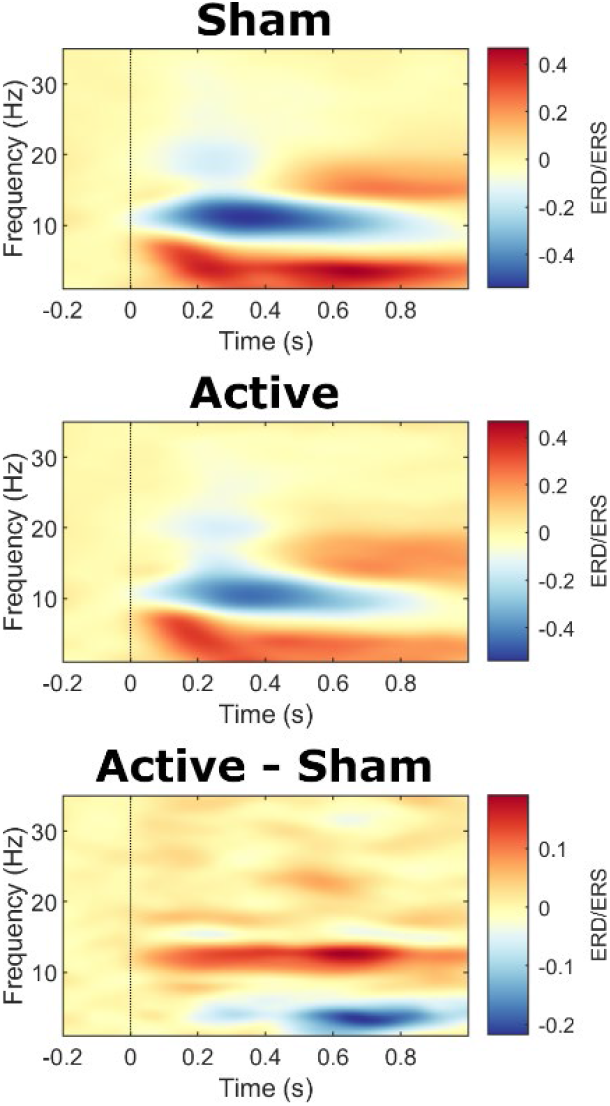
Time-frequency power during the 3-back task post-tDCS. EEG power shown for the average of frontal channels Afz and Fz for Sham and Active conditions, as well as the Active – Sham contrast.

**Figure 4.**
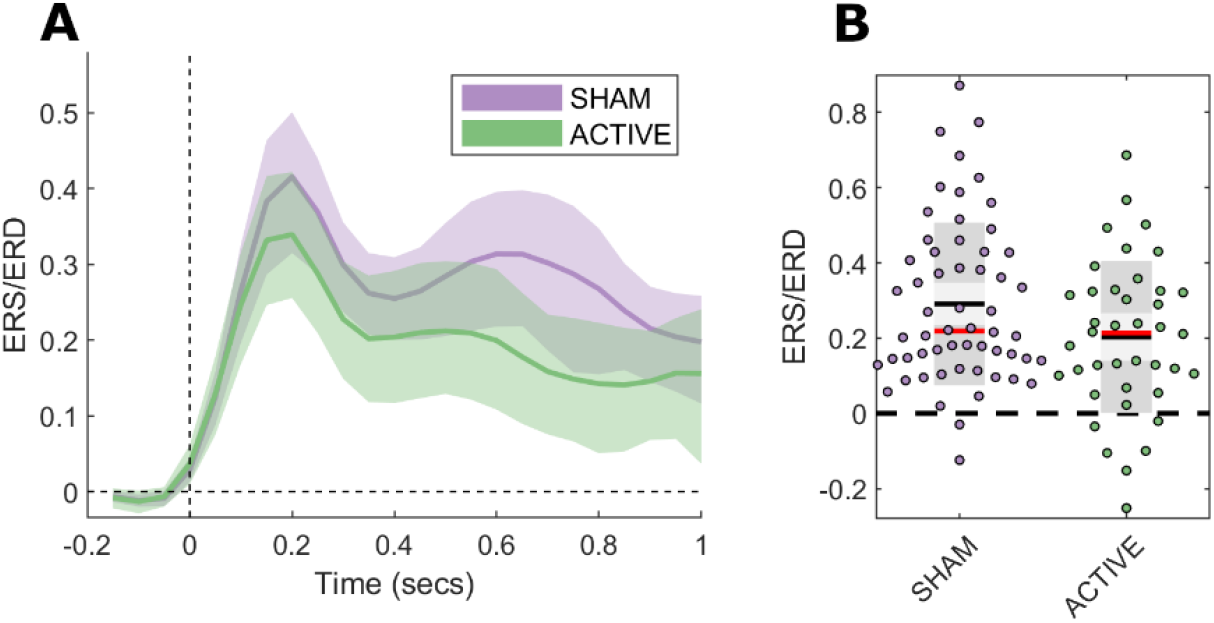
3-back task frontal theta event-related synchronisation post-tDCS. **A)** Average theta event-related synchronisation/desynchronization (ERS/D) over time displayed with bootstrapped 95% confidence intervals. **B)** Scatterplots of theta ERS/D obtained from frontal EEG channels AFz and Fz, 100 – 1000 ms post-stimulus onset. Black lines show the mean, red lines show the median, light grey shaded boxes indicate the 95% confidence interval, and dark grey regions indicate the standard deviation.

**Figure 5.**
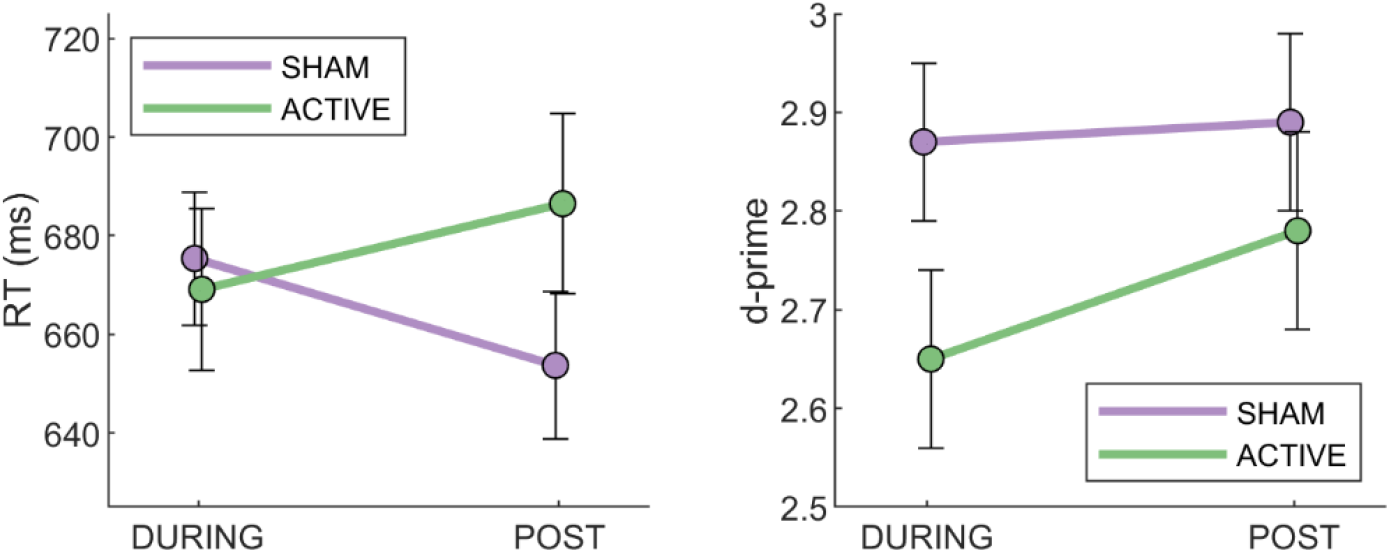
3-back task working memory performance outcomes. Line graphs show estimated marginal means for response time (RT) and accuracy (d-prime) during-, and post-tDCS. Baseline (pre-tDCS) performance was added as a covariate to account for individual differences in working memory. Error bars denote the standard error of the mean (SEM).

### Correlation

There were no significant correlations between event related spectra and working memory performance outcomes during- and post-tDCS following correction for the false discovery rate (all p > 0.05; Appendix Table A2).

### Blinding

Upon completion of the data collection phase of the experiment most participants guessed they had received active stimulation, 36/59 (61%) in the Sham group and 28/40 (70%) in the Active group. A Pearson’s chi-square test found no significant difference in guesses between groups (χ^2^ = 0.84, *p* = 0.359).

## Discussion

We investigated the concurrent and immediate aftereffects of tDCS on task-related spectral EEG activity during a letter 3-back working memory task. The Active group showed attenuated frontal theta ERS following tDCS, while no neurophysiological group differences were observed during tDCS. Behavioural measures showed a practice effect, with participants responding faster after than during the intervention, which was attenuated in participants receiving Active stimulation (1 mA or 2 mA) compared to those in the Sham tDCS group (0 mA, 0.016 mA, 0.034 mA).

Task-related frontal theta ERS was selectively affected out of all investigated event-related EEG outcomes following tDCS. Theta ERS plays a role in integrating and coordinating brain activity, including organisation of the multiple concurrent cognitive processes required for working memory (Sauseng, Griesmayr, Freunberger, & Klimesch, 2010). Further, theta is associated with higher-order executive functioning (A. Baddeley, 2003; Klimesch, Schack, & Sauseng, 2005; Sauseng, Klimesch, Schabus, & Doppelmayr, 2005), including continuous maintenance and manipulation processes required for the *n*-back task (Maurer et al., 2015; Pesonen et al., 2007; Sauseng et al., 2010), as well as cognitive control (Cavanagh & Frank, 2014; Duprez et al., 2020). The time course of frontal theta ERS reflects these functions, arising immediately after stimulus presentation and remaining elevated for at least a second to facilitate maintenance operations (Pesonen et al., 2007). An increase in theta power is therefore expected to be accompanied by improved working memory performance (Popov et al., 2018). Conversely, a decrease in task-related theta ERS should be associated with poorer performance (Missonnier et al., 2006), although the present study was unable to identify a significant correlation between theta activity and either d-prime or response latencies. Nevertheless, our findings could suggest that tDCS interferes with maintenance and cognitive control processes, indexed by frontal theta, early in the cognitive cascade leading to a correct response on the *n*-back task, although the strength of this interference did not rise to a level that produced noticeable impairments in performance. Alternatively, theta ERS has been shown to positively correlate with cognitive load, i.e., the number of letters ‘n’ to be recalled during the n-back task (Gevins & Smith, 2000; Gevins, Smith, McEvoy, & Yu, 1997; Jensen & Tesche, 2002; Pesonen et al., 2007). One might suggest that a decrease in theta following tDCS signifies an improvement in the efficiency of working memory processes and a relative reduction in cognitive load. However, attenuated practice effects in the active may group argue against this positive interpretation of reduced theta findings. Importantly, as theta could not be assessed during tDCS, it is unclear whether this disruption occurred during stimulation, or was the result of a neuromodulatory aftereffect. fMRI evidence suggests that 20 minutes of prefrontal tDCS can perturb cortical dynamics within the first 6 minutes of stimulation, and these effects remained stable for up to three days (Tu et al., 2021). It is therefore possible that a similar time-course might be present for theta activity, although this is obscured due to electrical artefacts in the EEG.

Similar effects of tDCS on theta activity have been reported by Powell, Boonstra, Martin, Loo, and Breakspear (2014) in a tDCS study of patients with an affective disorder. They showed a reduction in frontal theta during the retention phase of a verbal working memory task following tDCS, though this was not accompanied by significant behavioural effects between active and sham stimulation conditions. Conversely, Hoy et al. (2013) reported the opposite, finding enhanced frontal theta ERS in the active tDCS conditions (1 mA and 2 mA) relative to sham, although a significant effect was only observed during the 2-back but not the 3-back task. Similarly, anodal tDCS has been shown to increase theta ERS during a 2-back task relative to cathodal stimulation, but not compared to sham stimulation (Zaehle et al., 2011). Despite these studies indicating greater theta ERS following tDCS, others have not observed significant theta band differences between conditions in healthy (Hill et al., 2018; Ikeda et al., 2019; Murphy et al., 2020; Splittgerber et al., 2020) or patient populations (Hoy et al., 2015).

It is unclear which methodological difference/s may explain the disagreement between these studies. Potentially relevant factors include the timing of stimulation relative to task delivery, the tDCS montage, and the task load, though none of these sufficiently explains all discrepancies. The present study delivered the 3-back task during stimulation, which improves tDCS effects (Martin, Liu, Alonzo, Green, & Loo, 2014), possibly by modulating activity in task-relevant regions via functional specificity (Bikson & Rahman, 2013). Despite using a similar protocol, delivering tDCS during a task, Splittgerber et al. (2020) did not observe theta effects. Our study and Powell et al. (2014) used a bilateral F3/F4 or F3/F8 montage, both finding reduced theta following tDCS. Computational modelling suggests that F3/F4-F8 montages result in more right lateralised electric-fields, including the right DLPFC and orbitofrontal cortex, compared to montages in which the cathode is placed over the contralateral supraorbital region (Bai, Dokos, Ho, & Loo, 2014). These regions are consistently implicated in spatial and object processing (Mesulam, 2000; Owen et al., 2005; Reuter-Lorenz et al., 2000), and typically show significantly increased theta coherence during visuospatial working memory tasks at high cognitive loads (Muthukrishnan, Soni, & Sharma, 2020). Speculatively bilateral montages may therefore disrupt theta coherence within these regions to a greater degree than other commonly used electrode placements, however, further research is needed to confirm this hypothesis. Finally, it is possible that the use of the 3-back task, which entails a higher working memory load compared to the 2-back version of the task, may interact differentially with tDCS stimulation to produce reductions in theta. Similar effects have been reported in other cognitive domains, showing that tDCS reduced visual sustained attention at high but not low/medium cognitive loads (Roe et al., 2016). Likewise, results from a visuomotor tracking task found cognitive enhancing effects of tDCS at moderate, but not high difficulty levels (Kwon, Kang, Son, & Lee, 2015). Our results may therefore agree with a growing body of literature suggesting that task load is an important factor for consideration in assessing tDCS outcomes (de Almeida, Pope, & Hansen, 2020; Gill, Shah-Basak, & Hamilton, 2015; Meiron & Lavidor, 2013; Sánchez, Masip, & Gómez-Ariza, 2020).

Previous analyses of the present dataset found no significant differences for working memory accuracy and response times between the five conditions that were assessed (Nikolin et al., 2018). Categorisation of participants into broader Sham and Active groups, which increased statistical power, now shows relatively prolonged response latencies in Active (1 mA and 2 mA) compared to Sham stimulation (0 mA, 0.016 mA, 0.034 mA) for the post-tDCS aftereffect period (Cohen’s *d* = - 0.28). The observed interaction effect suggests that tDCS may have interfered with *n*-back task practice effects. These findings run counter to previous meta-analyses suggesting small-to-moderate benefits of tDCS to working memory performance (Brunoni & Vanderhasselt, 2014; Hill et al., 2016; Lee et al., 2021), and more generally improved response times for cognitive tasks following anodal prefrontal stimulation in healthy individuals (Dedoncker et al., 2016). Our results raise questions regarding the utility of tDCS to enhance *n*-back task performance, in agreement with meta-analytic findings showing no evidential value of tDCS in working memory studies (Medina & Cason, 2017). Considering the unexpected direction of our findings, a possible interference in practice effects following tDCS rather than an enhancement of cognition, further independent replication by similarly large, adequately powered, studies capable of detecting small effects is needed.

No significant differences were observed between Active and Sham groups for concurrent EEG measures obtained during tDCS. There are several plausible interpretations of this outcome. Firstly, the only significant effect observed post-tDCS occurred in the theta frequency band, which could not be assessed during stimulation due to excessive artefacts. Secondly, there is evidence to suggest that the aftereffects of tDCS are larger than the concurrent effects, and so may be statistically easier to detect (Jamil et al., 2020). Thirdly, qualitatively larger interindividual variability was observed during-tDCS. The resulting heterogeneity in outcomes, a common feature of tDCS (Li, Uehara, & Hanakawa, 2015), reduces statistical power to identify group differences. Finally, some stimulation artefacts may still have been present in the EEG data acquired during tDCS despite cleaning efforts, thereby obscuring the effects of stimulation (Boonstra, Nikolin, Meisener, Martin, & Loo, 2016). Although simple temporal and spatial filtering strategies may attenuate some tDCS-induced noise (Mancini et al., 2015; Marghi et al., 2015), they appear to be insufficient to eradicate all inherent and non-inherent physiological and stimulator artefacts. These pose a unique non-stationary problem that can only be removed with significant effort (Gebodh et al., 2019).

An advantage of the present study compared to prior investigations of similar outcomes is the large sample size afforded by analysis of combined data collected over two experiments. This allowed for the detection of moderate frontal theta ERS (Cohen’s d = -0.42) and small response time (Cohen’s d = -0.28) effects of stimulation. Interestingly, our findings indicate that neurophysiological measures may be more sensitive to the effects of tDCS than cognitive performance outcomes. Indeed, similar views have been expressed using event-related potentials obtained during the *n*-back task (Hill, Rogasch, Fitzgerald, & Hoy, 2019; Keeser et al., 2011), including previously reported analyses of the present dataset (Nikolin et al., 2018). These findings suggest a greater role for EEG, and more broadly neuroimaging techniques, including functional magnetic resonance imaging (Fischell, Ross, Deng, Salmeron, & Stein, 2020; Lin et al., 2019; Vaqué-Alcázar et al., 2020) and functional near infra-red spectroscopy (Schommartz, Dix, Passow, & Li, 2021), for investigations of the neuromodulatory effects of tDCS on cognitive processes. In the absence of behavioural changes following tDCS, these techniques may be used instead to gain insights into the mechanisms of action of tDCS, and thereby optimise stimulation parameters to achieve larger and more consistent cognitive effects.

## Limitations

The data presented in the current study was collected in two separate experiments and so randomisation was not possible to all five conditions at once, potentially introducing a sampling bias. However, the inclusion criteria and method of recruitment were purposefully kept consistent in both experiments, such that participants were similar in demographic and working memory outcomes at baseline. Additionally, identical working memory performance thresholds were used in both experiments during randomised stratification of participants to stimulation conditions. Although the combination of these datasets introduces heterogeneity due to inconsistent stimulation parameters, in doing so it may improve the generalisability of findings. A consensus is yet to emerge for the optimal tDCS current intensity necessary to achieve cognitive enhancing effects of tDCS. As a result, the vast majority of studies select dosage parameters ranging between 1 – 2 mA (Hill et al., 2016), as was used in the present study. Our results may thus be broadly reflective of tDCS working memory effects obtained using commonly employed current intensities within the field. We were unable to measure frontal theta during tDCS due to significant electrical artefacts introduced to the EEG signal in this frequency band. Hence, we cannot determine whether differences in theta could have emerged during this time-point, preceding response time effects and potentially suggesting a causal link. Lastly, the interpretation of tDCS causing frequency specific alterations in the theta band should be made with caution. Although previous research has labelled similar findings as an effect on ‘oscillatory activity’ (Heinrichs-Graham, McDermott, Mills, Coolidge, & Wilson, 2017; Hill et al., 2018; Ikeda, Takahashi, Hiraishi, Saito, & Kikuchi, 2019; Murphy et al., 2020; Zaehle, Sandmann, Thorne, Jäncke, & Herrmann, 2011), changes in spectral power following stimulation may instead occur due to generalised shifts in the balance of excitation and inhibition within the brain. In such instances, signal fitting algorithms may be used to distinguish oscillatory and aperiodic components in the power spectrum (Donoghue et al., 2020). While this technique was recently applied to event-related data (Virtue-Griffiths et al., 2022), testing whether it can accurately distinguish rapid changes in event-related theta from changes in the slope of aperiodic activity is beyond the scope of this study.

## Conclusions

Our findings suggest that tDCS may interfere with working memory processes, particularly maintenance and cognitive control as measured by frontal theta ERS. This may have attenuated practice effects, resulting in a relative lack of improvement in response latencies post-tDCS as compared to participants receiving Sham stimulation. The analysis of task-related spectral EEG changes provides insights into mechanisms by which tDCS affects working memory, in particular, why tDCS may not always be useful in augmenting working memory in healthy volunteers.

## Acknowledgements

Dr Nikolin was supported by an Australian Postgraduate Award during data collection. Dr Martin and Dr Boonstra both were the recipients of a NARSAD Young Investigator Grant. The authors have no conflicts of interest to declare.

## Appendix

**Appendix Figure A1.**
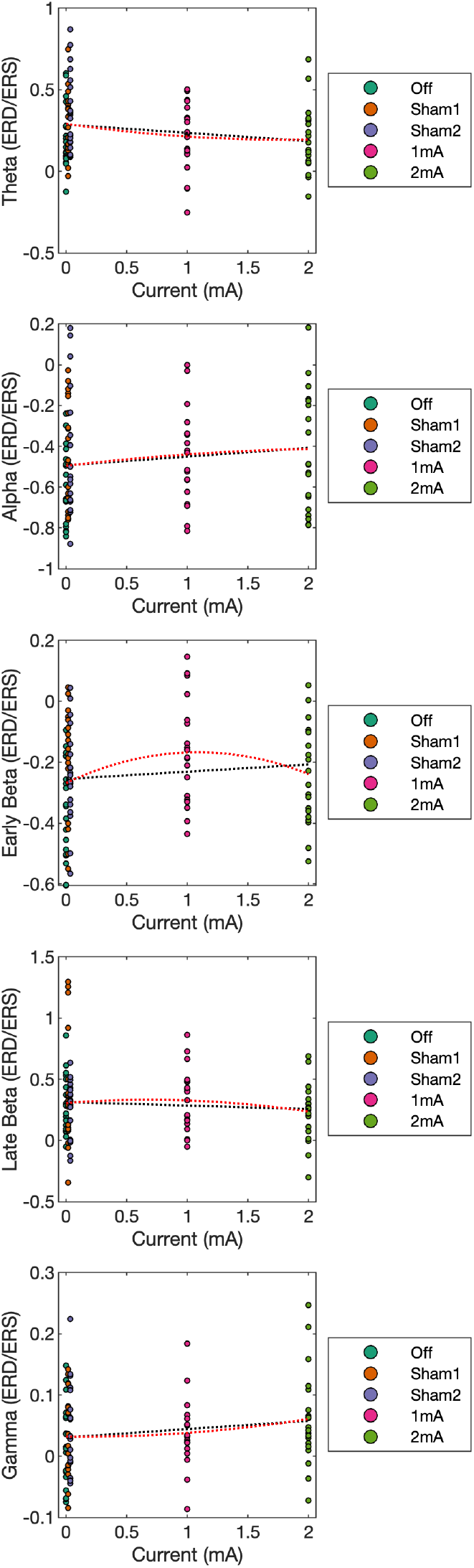
Scatterplots of simple univariate linear regression for during-tDCS time-frequency outcomes. Time-frequency measures were set as the dependent variable, with current intensity (mA) included as the independent variable. Black dotted lines show the line of best fit for 1^st^ order linear regressions, and red dotted lines show the same for 2^nd^ order linear regressions.

**Appendix Table A1.**
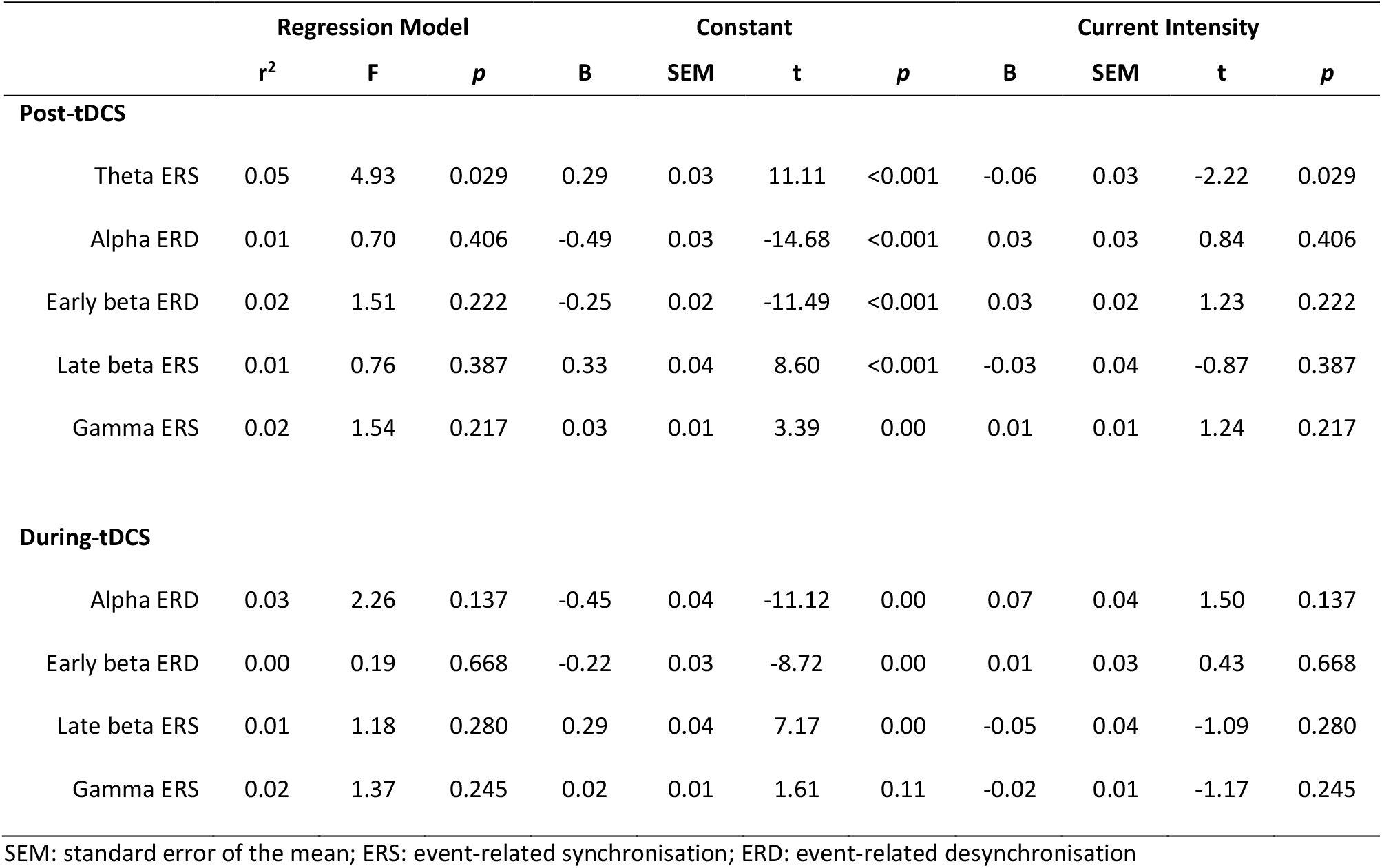
Simple univariate linear regression outcomes

|  | Regression Model |  |  | Constant |  |  |  | Current Intensity |  |  |  |
| --- | --- | --- | --- | --- | --- | --- | --- | --- | --- | --- | --- |
|  | *r*<sup>2</sup> | F | *p* | B | SEM | t | *p* | B | SEM | t | *p* |
| **Post-tDCS** |  |  |  |  |  |  |  |  |  |  |  |
| Theta ERS | 0.05 | 4.93 | 0.029 | 0.29 | 0.03 | 11.11 | <0.001 | -0.06 | 0.03 | -2.22 | 0.029 |
| Alpha ERD | 0.01 | 0.70 | 0.406 | -0.49 | 0.03 | -14.68 | <0.001 | 0.03 | 0.03 | 0.84 | 0.406 |
| Early beta ERD | 0.02 | 1.51 | 0.222 | -0.25 | 0.02 | -11.49 | <0.001 | 0.03 | 0.02 | 1.23 | 0.222 |
| Late beta ERS | 0.01 | 0.76 | 0.387 | 0.33 | 0.04 | 8.60 | <0.001 | -0.03 | 0.04 | -0.87 | 0.387 |
| Gamma ERS | 0.02 | 1.54 | 0.217 | 0.03 | 0.01 | 3.39 | 0.00 | 0.01 | 0.01 | 1.24 | 0.217 |
| **During-tDCS** |  |  |  |  |  |  |  |  |  |  |  |
| Alpha ERD | 0.03 | 2.26 | 0.137 | -0.45 | 0.04 | -11.12 | 0.00 | 0.07 | 0.04 | 1.50 | 0.137 |
| Early beta ERD | 0.00 | 0.19 | 0.668 | -0.22 | 0.03 | -8.72 | 0.00 | 0.01 | 0.03 | 0.43 | 0.668 |
| Late beta ERS | 0.01 | 1.18 | 0.280 | 0.29 | 0.04 | 7.17 | 0.00 | -0.05 | 0.04 | -1.09 | 0.280 |
| Gamma ERS | 0.02 | 1.37 | 0.245 | 0.02 | 0.01 | 1.61 | 0.11 | -0.02 | 0.01 | -1.17 | 0.245 |

SEM: standard error of the mean; ERS: event-related synchronisation; ERD: event-related desynchronisation

**Appendix Figure A2.**
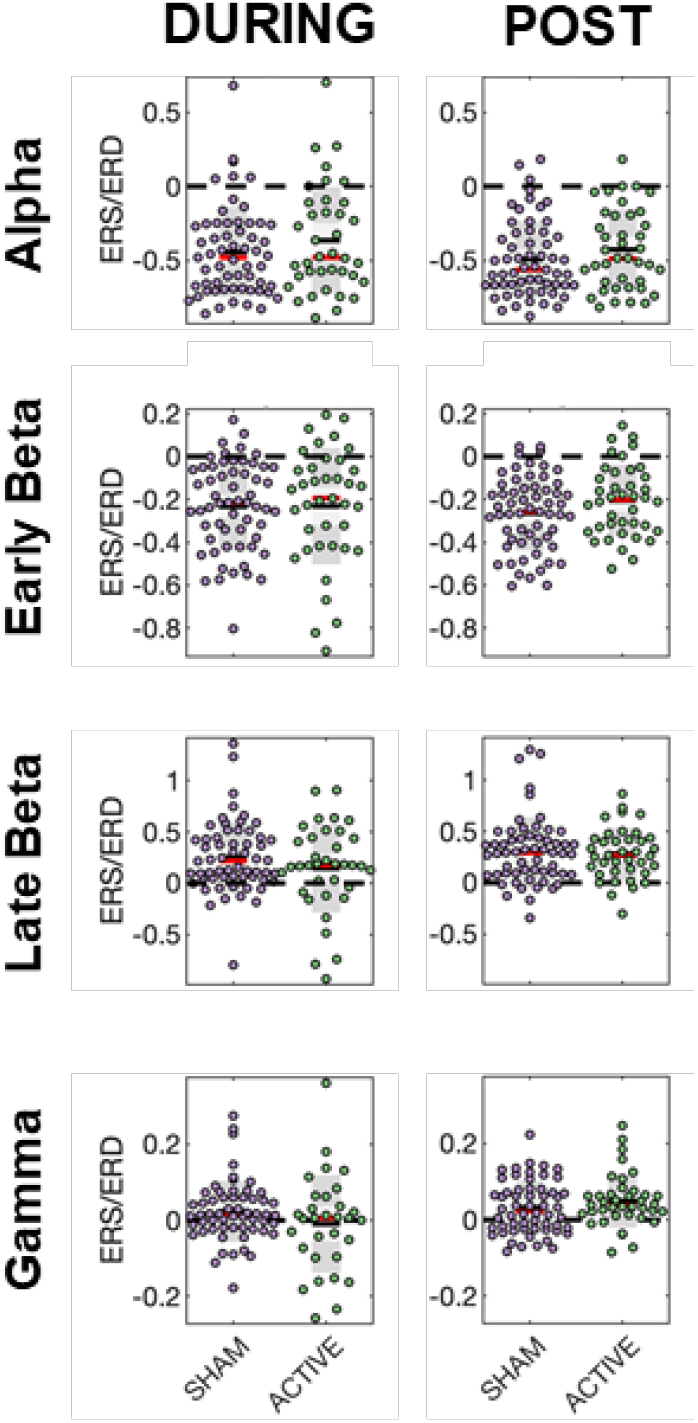
Scatterplots of other frequency band results.

**Appendix Table A2.** Pearson correlations comparing event-related time-frequency power spectra and working memory performance measures. Correlations were corrected for multiple comparisons using the false discovery rate.

|  | Response Time |  |  | D-prime |  |  |
| --- | --- | --- | --- | --- | --- | --- |
|  | *r* | *p* | FDR<br>*p* | *r* | *p* | FDR<br>*p* |
| **Post-tDCS** |  |  |  |  |  |  |
| Theta ERS | -0.06 | 0.553 | 0.664 | -0.08 | 0.438 | 0.606 |
| Alpha ERD | -0.08 | 0.428 | 0.606 | -0.17 | 0.093 | 0.422 |
| Early beta ERD | -0.15 | 0.132 | 0.422 | -0.11 | 0.274 | 0.529 |
| Late beta ERS | -0.14 | 0.164 | 0.422 | -0.05 | 0.611 | 0.687 |
| Gamma ERS | 0.11 | 0.294 | 0.529 | -0.14 | 0.163 | 0.422 |
| **During-tDCS** |  |  |  |  |  |  |
| Alpha ERD | -0.10 | 0.356 | 0.583 | -0.01 | 0.927 | 0.964 |
| Early beta ERD | -0.11 | 0.287 | 0.529 | 0.01 | 0.964 | 0.964 |
| Late beta ERS | -0.07 | 0.485 | 0.624 | 0.15 | 0.153 | 0.422 |
| Gamma ERS | -0.20 | 0.063 | 0.422 | 0.27 | 0.010 | 0.180 |

FDR: false discovery rate; RT: response time; tDCS: transcranial direct current stimulation;
ERS: event-related synchronisation; ERD: event-related desynchronisation

## Sensitivity Analyses

Our previous analyses indicated that the Sham2 group may have biological effects (Nikolin et al., 2018). However, Sham2 amplitudes were not significantly different to Sham1, and indeed were remarkably similar (Sham1 P3 = -0.59 µV; Sham2 P3 = -0.57 µV; p = 0.95, Hedges’ g = 0.02. Considering concerns that this group may alter brain activity, we re-ran analyses excluding participants from each group, similar to a condition-level leave-one-out analysis, to assess the impact of this condition on our overall findings.

Condition-level leave-one-out behavioural analyses resulted in similar response time outcomes. The main effect of Time remained significant, indicating shorter latencies post-tDCS relative to during-tDCS, except following exclusion of the Off group in which it was at trend level (*F* = 3.4, *p* = 0.07, η^2^ = 0.04). The main effect of Condition was non-significant following all exclusions. The Time × Condition interaction remained significant except following exclusion of the 1 mA group (*F* = 1.9, *p* = 0.18, η^2^ = 0.02).

Likewise, d-prime analysis revealed no significant main effect of Time, and no Time × Condition interaction following each group exclusion. The main effect of Condition narrowly attained the significance threshold following exclusion of Sham2 participants (*F* = 4.0, *p* = 0.05, η^2^ = 0.05), indicating reduced working memory accuracy in the Active group.

The MANOVAs for EEG outcomes were no longer significant in the time-period post tDCS following exclusion of Off (*p* = 0.10), 1mA (*p* = 0.09), and 2mA (*p* = 0.13) conditions. Only exclusion of the Sham1 group resulted in significant post-tDCS EEG findings (*p* = 0.02). For the time-period during tDCS, all exclusions resulted in non-significant MANOVAs similar to results obtained using the full sample (Off *p* = 0.66; Sham1 *p* = 0.07; Sham2 *p* = 0.60; 1 mA *p* = 0.16; 2 mA *p* = 0.21).

